# Picobiinae mites (Acariformes: Syringophilidae) parasitising the Starlings (Passeriformes: Sturnidae) in the Afrotropical region

**DOI:** 10.1101/2024.07.09.602639

**Authors:** Milena Patan, Maciej Skoracki, Iva Marcisova, Martin Hromada, Bozena Sikora

**Affiliations:** Department of Animal Morphology, Faculty of Biology, Adam Mickiewicz University, Uniwersytetu Poznańskiego 6, 61-614 Poznan, Poland; Laboratory and Museum of Evolutionary Ecology, Department of Ecology, Faculty of Humanities and Natural Sciences, University of Prešov, 080 01 Prešov, Slovakia; Faculty of Biological Sciences, University of Zielona Góra, Prof. Z. Szafrana 1, 65-516, Zielona Góra, Poland

**Keywords:** Acari, Africa, birds, ectoparasites, taxonomy

## Abstract

In the present paper, we continue our studies on quill mites of the family Syringophilidae parasitising birds of the family Sturnidae. Herein, we describe a new species, *Picobia wisniewskii* **sp. nov**., collected from the red-winged starling *Onychognathus morio* (Linnaeus) in Tanzania. Additionally, we provide an emended diagnosis and new host records for *Picobia lamprotornis* Klimovicova et al., 2014 and *Picobia sturni* Skoracki *et al*. 2004.

## INTRODUCTION

The Afrotropical region is a biodiversity hotspot with a rich and diverse bird community (Fjeldså *et al*. 2020). Among the most evolutionarily successful are the Passeriformes, a dominant order that displays various adaptations and morphological forms. Passerines, commonly known as perching birds or songbirds, represent the most diverse and widespread order of birds in the world, and the Afrotropical region is no exception. Passerine birds found in this region exhibit extraordinary diversity in size, colouration, and habitat preferences, reflecting the diverse ecological niches they occupy. From dense rainforests to arid savannahs, these birds have adapted to virtually all available habitats in this vast area (Winkler *et al*. 2020).

Among passerine birds, the representatives of the family Sturnidae, which includes starlings, mynas, and rhabdornises (Lovette & Rubenstein 2007; Clements *et al*. 2023), are known for their often highly social behaviour and forming large flocks (Winkler *et al*. 2020). The world fauna of the starlings comprises approximately 125 species grouped in 36 genera. Among these, 48 species in 13 genera belong to the Afrotropical avifauna (Lepage *et al*. 2014). According to Fjeldså *et al*. (2020), lineages of African starlings diverged from the sister Asian savanna starling clade relatively recently, around 10 Mya.

The adaptation of starlings to different ecological niches provides an interesting background for studying their relationships with parasitic prostigmatan mites.

Representatives from most families of prostigmatan mites permanently associated with birds, i.e., Cheyletidae, Ereynetidae, and Syringophilidae, have been identified on starlings in the Afrotropical region. However, these families have yet to undergo a comprehensive revision. For instance, the family Cheyletidae is represented by only two species associated with two starling species (Fain 1972, 1981; Fain & Bochkov 2002), and Ereynetidae by merely three species associated with five sturnid host species (Fain 1963, 1964, 1971). The family Syringophilidae recorded on Afrotropical starlings until now comprises three species in three genera, i.e., *Syringophiloidus soponai* Skoracki *et al*., 2024, *Syringophilopsis parasturni* Skoracki *et al*., 2024, and *Picobia lamprotornis* Klimovicova *et al*., 2004 (see Skoracki *et al*. 2024).

This paper reports new observations about quill mites of the subfamily Picobiinae parasitising African starlings. Herein, we describe a new species *Picobia wisniewskii* **sp. nov**., and provide an emended diagnosis and new host records for *P. lamprotornis* Klimovicova *et al*., 2014, and new host record for *P. sturni* Skoracki *et al*., 2004.

## MATERIAL AND METHODS

The mite specimens used in this study were collected from dried bird skins housed in the ornithological collections of the Bavarian State Collection of Zoology (SNSB - Zoologische Staatssammlung München), Munich, Germany. The collection methodology followed Skoracki (2011). We examined the quills of approximately ten contour feathers in the proximity of the cloaca region for each bird. Prior to mounting, mites were removed from feathers, softened, and cleared in Nesbitt’s solution at 40°C for three days. Mite specimens were identified using a light microscope (ZEISS Axioscope2™; Carl Zeiss AG, Jena, Germany) with differential interference contrast (DIC) illumination. Drawings were created with a camera lucida.

The description of idiosomal setation adheres to Grandjean (1939) as adapted for Prostigmata by Kethley (1990), while the nomenclature of leg setae follows Grandjean (1944). The morphological terminology is based on Skoracki (2011) and Skoracki *et al*. (2016). All measurements are in micrometres (μm), with ranges for paratypes in parentheses given after data for a holotype.

The scientific names of birds follow Clements *et al*. (2023). The collected mite material is now housed at the Adam Mickiewicz University, Department of Animal Morphology, Poznań, Poland (AMU) and the Bavarian State Collection of Zoology, Arthropoda Varia, Munich, Germany (SNSB-ZSM).

## RESULTS

**Family Syringophilidae Lavoipierre, 1953**

**Subfamily Picobiinae Johnston & Kethley, 1973**

**Genus *Picobia* Haller, 1887**

### *Picobia wisniewskii* Patan, Skoracki & Marcisova sp. nov

(Figures 1 and 2)

**Figure 1.**
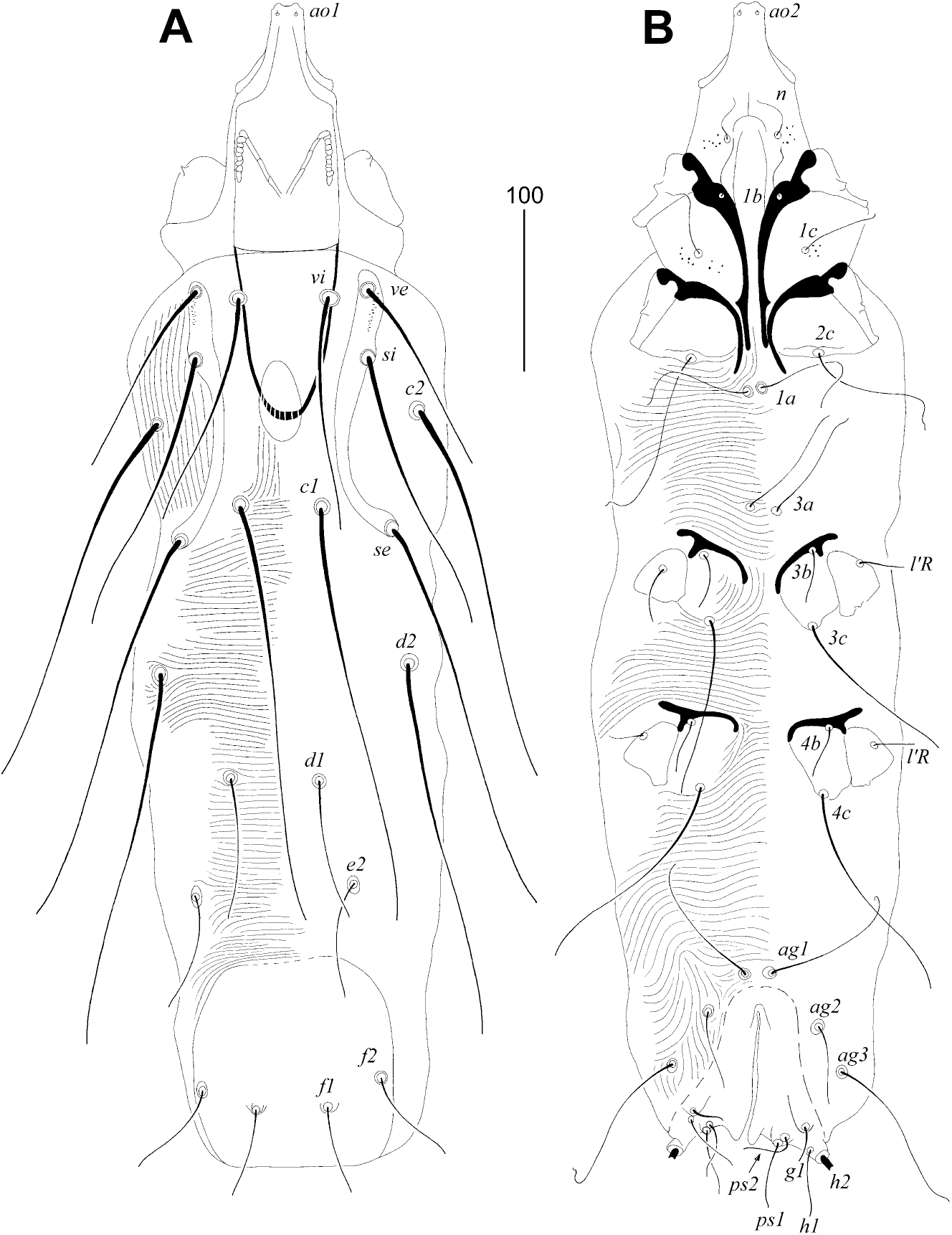
*Picobia wisniewskii* Patan, Skoracki & Marcisova sp. nov., female: (A) dorsal view; (B) ventral view.

**Figure 2.**
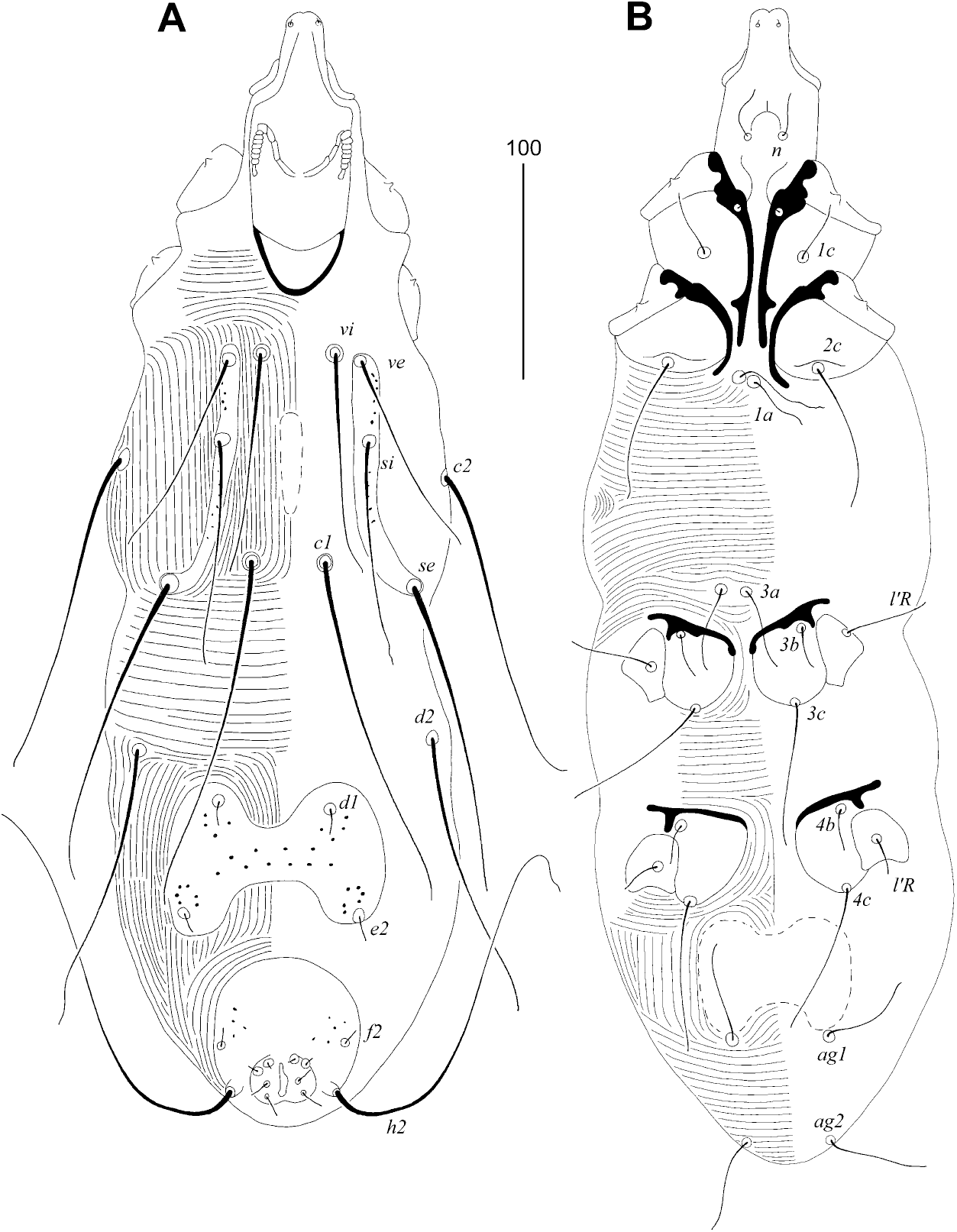
*Picobia wisniewskii* Patan, Skoracki & Marcisova sp. nov., male: (A) dorsal view; (B) ventral view.

#### Non-physogastric female (holotype and two paratypes)

Total body length 635. Gnathosoma. Hypostomal apex rounded, with pair of small shoulders. Infracapitulum sparsely punctate. Each medial branch of V-shaped peritremes with four chambers, each lateral branch with nine clearly visible chambers. Stylophore 240 (235) long, exposed portion of stylophore (stylophoral shield) apunctate 155 (150) long. Movable cheliceral digit 200 (200–205) long. Idiosoma. Propodontal shield divided into three sclerites, i.e., two narrow lateral sclerites bearing bases of setae *ve, si* and *se*, punctate between bases of setae *ve* and *si*, and small, oval, apunctate medial sclerite. Setae *vi* and *ve* situated at same transverse level.

Setae *c1* situated anterior to level of setae *se*. Propodontal setae *vi, ve* and *si* beaded, other propodontal and hysteronotal setae lightly ornamented. Setae *d1* situated equidistant between setae *d2* and *e2*. Pygidial shield well developed, apunctate. Setae *f1* slightly (1.1–1.3 times) longer than *h1*. Alveoli of setae *3a* not coalesced. Agenital setae *ag1* situated antero-medial to level of setae *ag2*. Genital plate present, apunctate. Genital setae situated on small, blunt-ended genital lobes. Pseudanal setae *ps1* 1.3–1.5 times longer than *ps2*. All coxal fields well developed, fields I punctate, II–IV apunctate. Legs. Antaxial and paraxial members of claws equal in size and shape. Lengths of setae: *vi* (135–140), *si* 175 (165), *se* 235 (250), *c1* 260 (255–275), *c2* 235 (230–240), *d1* 90 (85–90), *d2* 230 (230–240), *e2* 70 (70–90), *f1* 45 (40– 45), *f2* 55 (50–60), *h1* 35 (35–55), *ps1* 45 (40–45), *ps2* 35 (30), *g1* 25 (25), *ag1* 80 (75–85), *ag2* 40 (40–50), *ag3* 85 (85–105), *l’RIII* 35 (30–35), *l’RIV* 25 (25), *3b* 25 (25), *3c* 105 (125), *4b* 30 (25–30), *4c* 125.

#### Physogastric female

Features as in non-physogastric form except body worm-shaped outline.

#### Male

Total body length is 400–460 in two paratypes. Gnathosoma. Hypostomal apex rounded. Infracapitulum apunctate. Each medial branch of M-shaped peritremes with four chambers, each lateral branch with eight clearly visible chambers. Stylophore 110–120 long; exposed portion of stylophore apunctate, 90–105 long. Movable cheliceral digit 80 long.

Idiosoma. Propodontal shield divided into two narrow and sparsely punctate lateral sclerites, bearing bases of setae *ve, si, se*, and small narrow apunctate medial sclerite. Setae *vi* and *ve* situated at same transverse level. Setae *vi, ve* and *si* lightly beaded and subequal in length.

Setae *c1* situated anterior to level of setae *se*. Hysteronontal shield entire, not fused to pygidial shield, punctate, bearing bases of setae *d1* and *e2*, distinctly cleft on anterior and posterior margins. Setae *d1* situated equidistant between setae *d2* and *e2*. Pygidial shield well developed, rounded on anterior margin, sparsely punctate. Alveoli of setae *3a* not coalesced. Agenital plate entire, weakly sclerotised, apunctate, bearing bases of setae *ag1* on posterior margin. All coxal fields well developed and apunctate. Lengths of setae: *vi* 100–105, *ve* 90– 105, *si* 105–110, *se* 125–145, *c1* 165, *c2* 150, *d1* 10, *d2* 120–130, *e2* 10, *f2* 10, *h2* 145–205,*ag1* 30–55, *ag2* 25–35, *3b* 20, *3c* 65, *4b* 15–20, *4c* 60.

#### Type material

Female holotype (non-physogastric form) and paratypes: two females non-physogastric form, three females physogastric form, and two males (reg. no. AMU-MS 21- 1012-34) from feathers of the red-winged starling *Onychognathus morio* (Linnaeus) (host reg. no. SNSB-ZSM 64.717; female); TANZANIA: Morogoro District, 6 May 1962, alt. 1,500–1,800 m. a.s.l., coll. Th. Andersen. One female physogastric form (reg. no. AMU-MS 21-1012-035) from same host species (host reg. no. SNSB-ZSM 64.717; female) and locality.

#### Type material deposition

Holotype and most of paratypes are deposited in the SNSB-ZSM, except one female non-physogastric form, one female physogastric form and one male in the AMU.

#### Differential diagnosis

*Picobia wisniewskii* **sp. nov**. is morphologically similar to *Picobia lamprotornis* by the presence of short setae *e2* (shorter than 110) and differs from it by the presence of the following features: in females of *P. wisniewskii*, the length of stylophore is 235–240; the medial sclerite of the propodontal shield is punctate; the lengths of setae *c2* and *d2* are subequal 230–240; in males, the agenital plate is entire. In females of *P. lamprotornis*, the length of stylophore is 195–200; the medial sclerite of propodontal shield is apunctate; the lengths of setae *c2* and *d2* are 175–180 and 165–190, respectively; in males, the agenital plate is divided into two small and oval plates.

#### Etymology

This species is named in honour of the Polish acarologist Prof. Jerzy Wiśniewski from the University of Life Sciences in Poznań.

### *Picobia lamprotornis* Klimovicova, Skoracki, Wamiti & Hromada, 2014

Emended description

#### Female

Hypostomal apex rounded, with pair of small shoulders. Each medial and lateral branch of peritremes with four and eight chambers, respectively. Propodonotal shield divided into two narrow lateral shields and small oval and punctate medial shield. Pygidial shield sparsely punctate. Setae *ag1* situated antero-medial to setae *ag2*. Genital plate present, apunctate. Genital setae situated on small, rounded genital lobes. Both tarsal claws subequal in size and shape. Measurements. Stylophore 195 (195–200); lengths of setae: *ve* 90–95, *si* 150, *se* 190, *c1* 215–230, *c2* 175–180, *d2* 165–190, *e2* 95–100, *f1* 35, *f2* 50, *h1* 30–50, *ps1* 20–30, *ps2* 25, *ag1* 40–45, *ag2* 30–35, *ag3* 65–90, *3b* 30–35, *3c* 70–100.

#### Male

Total body length 455–480. Each medial branch of peritremes with three or four chambers, each lateral branch with seven or eight chambers. Propodonotal shield divided into two narrow lateral shields and single medial shield, all sclerites punctate. Hysteronotal shield entire, punctate. Pygidial shield apunctate. Two oval agenital plates present. Lengths of setae: *vi* 100–105, *ve* 85–90, *si* 110–120, *c1* 155–160, *c2* 120–130, *d2* 130, *e2* 5, *h2* 220, *ps1* 5, *ps2* 5, *g1* 5, *g2* 5, *ag1* 35–45, *ag2* 20, *l’RIII* 20, *tc’III–IV* 30–40, *tc”III–IV* 45, *3b* 25, *3c* 45–50.

#### Host and distribution

Mite species associated with Afrotropical starlings (Sturnidae): the superb starling *Lamprotornis superbus* Rueppell (type host) from Kenya (Klimovicova *et al*. 2014), the greater blue-eared glossy-starling *Lamprotornis chalybaeus* (Hemprich & Ehrenberg) from Tanzania and Kenya, the lesser blue-eared glossy-starling *Lamprotornis chloropterus* (Swainson) from Tanzania (Skoracki *et al*. 2024), the Abbott’s starling *Poeoptera femoralis* (Richmond) from Tanzania and the Kenrick’s starling *Poeoptera kenricki* (Shelley) from Tanzania (current study).

#### New material examined

Three females non-physogastric form, two females physogastric form, and three males (reg. no. AMU-MS 21-1012-042) from feathers of the Abbott’s starling *Poeoptera femoralis* (Richmond) (host reg. no. SNSB-ZSM 59.148; male); TANZANIA: Arusha Region, Arusha National Park, Mt Meru, 2,100 m. a.s.l., 16 November 1958, coll. v. Nagy.

Three females physogastric form and four males (reg. no. MS 21-1012-006) from contour feather quill of the Kenrick’s starling *Poeoptera kenricki* (Shelley) (host reg. no. SNSB-ZSM 60.1797; male); TANZANIA: Arusha Region, Meru District, Maji ya Chai, 1,600 m. a.s.l., 30 December 1959, coll. unknown.

### *Picobia sturni* Skoracki, Bochkov & Wauthy, 2004

To this time this species was recorded from the Eurasian starling *Sturnus vulgaris* Linnaeus in Poland, Moldova and Slovakia (Skoracki *et al*. 2004, 2010, 2016; Skoracki 2011), the spotless starling *Sturnus unicolor* Temminck in Morocco (Skoracki *et al*. 2016), and from the white-cheeked starling *Spodiopsar cineraceus* (Temminck) in Japan (Skoracki 2011).

Herein, we give a new record of this mite species from the wattled starling *Creatophora cinerea* (Meuschen) in Kenya.

#### New material examined

Two females (physogastric form) and two males (reg. no. AMU-MS 21-1012-040) from contour feather quill of the wattled starling *Creatophora cinerea* (Meuschen) (host reg. no. SNSB-ZSM 26.397; male); KENYA: Mount Elgon National Park, Mount Elgon steppe, 2,300 m. a.s.l., 8 February 1925, coll. S. Alinder.

## DISCUSSION

The genus *Picobia* is recognised for its exceptional taxonomic diversity. As a member of the *Picobia*-generic-group (see Skoracki *et al*. 2020), it currently encompasses 49 species, distinguishing itself by having a higher number of species compared to other picobiine genera, which do not surpass ten described taxa. This genus is predominantly associated with passerine birds (Passeriformes; clade Australaves), although a few species have also been recorded on woodpeckers (Piciformes; clade Afroaves). Thus, the genus *Picobia* is well established in both main clades of Telluraves (Ericson 2012; Jarvis *et al*. 2014). To date, *Picobia* representatives have been recorded in 24 passerine families (comprising both oscines and suboscines): Aegithalidae, Alaudidae, Corcoracidae, Corvidae, Estrildidae, Fringillidae, Furnariidae, Hirundinidae, Leiothrichidae, Muscicapidae, Nectariniidae, Panuridae, Paradisaeidae, Passeridae, Pellorneidae, Ploceidae, Pycnonotidae, Rhinocryptidae, Sturnidae, Sylviidae, Thamnophilidae, Troglodytidae, Turdidae (suborder Oscines), and Tyrannidae (suborder Suboscines). In addition to its broad host range, this genus has been recorded in all zoogeographical regions except Antarctica (Skoracki *et al*. 2016). Picobiine mites associated with starlings comprise only four species (including a new species described herein) recorded on 13 host species (Skoracki *et al*. 2024; current study). Notably, each zoogeographical region has its own set of mites parasitising starlings. In the Afrotropical region, three species have been identified on various starling species: *Picobia lamprotornis*, associated with three bird species of the genus *Lamprotornis* (*L. superbus, L. chalybaeus, L. chloropterus*) and two species of the genus *Poeoptera* (*P. femoralis* and *P. kenricki*) (Klimovicova *et al*. 2014; Skoracki *et al*. 2024; current study); *Picobia wisniewskii* associated with *Onychognathus morio*, and *Picobia sturni* parasitising in Africa on *Creatophora cinerea* (current study).

The wattled starling *Creatophora cinerea* is the only African starling that appears to show affinities with the Asian starlings; according to Fjeldså *et al*. (2020) it is a member of Eurasian savannah starling clade, including the genus *Sturnus* as a basal lineage. The rest of our African host assemblages are part of the clade sister to Eurasian savannah starlings - African red-winged starling lineage (*Onychognathus*) and African glossy starlings (*Lamprotornis* and *Poeoptera*) (Fjeldså *et al*. 2020).

In the Palaearctic region, only one species has been recorded to date, *Picobia sturni* Skoracki, Bochkov & Wauthy, 2004, associated with three host species: *Sturnus vulgaris, S. unicolor*, and *Spodiopsar cineraceus*, all members of Eurasian savannah clade according to Fjeldså *et al*. (2020). Finally, in the Oriental/Oceanian region, only one species has been recorded, *Picobia indonesiana* Skoracki & Glowska, 2008, parasitising three host species: *Aplonis panayensis, Enodes erythrophris*, and *Mino dumontii* (Skoracki & Glowska 2008), representing more basal Mainathinae sturnid radiation (Fjeldså *et al*. 2020).

## Acknowledgements

We express our gratitude to Dr. Markus Unsöld (Ornithological Section, Bavarian State Collection of Zoology, Munich, Germany (SNSB-ZSM)) for making available the samples of dry bird skins for the present study. We also thank all members of the Society - Freunde der Zoologischen Staatssammlung München e. V. - for their invaluable support during our research tenure at SNSB-ZSM. This study was supported by the AMU Excellence Initiative – Research University, grant no. 118/34/UAM/0056 (to M.P.) and by the Slovak Research and Development Agency under the contract APVV-22-0440 (to M.S., I.M. and M.H.) and the Agency of the Ministry of Education, Research and Sport of the Slovak Republic and Slovak Academy of Sciences 1/0876/21 (to M.S., I.M., M.H., and B.S).

